# Are oceanic fronts ecotones? Seasonal changes along the Subtropical Front show fronts as bacterioplankton transition zones but not diversity hotspots

**DOI:** 10.1101/109868

**Authors:** Sergio E. Morales, Moana Meyer, Kim Currie, Federico Baltar

## Abstract

Ecotones are regarded as diversity hotspots in terrestrial systems, but it is unknown if this “ecotone effect” occurs in the marine environment. Oceanic fronts are widespread mesoscale features, present in the boundary between different water masses, and are arguably the best potential examples of ecotones in the ocean. Here we performed the first seasonal study along an oceanic front, combining 16S rRNA gene sequencing coupled with a high spatial resolution analysis of the physical properties of the water masses. Using the Subtropical Frontal Zone off New Zealand we demonstrate that fronts delimit shifts in bacterioplankton community composition between water masses, but that the strength of this effect is seasonally dependent. While creating a transition zone where physicochemical parameters and bacterioplankton communities get mixed, this ecotone does not result in increased diversity. Thus unlike terrestrial ecotones, oceanic ecotones like fronts are boundaries but not hotspot of bacterioplankton diversity in the ocean.

---

Ecotones are boundaries, or transition zones, between ecological communities, ecosystems, or ecological regions usually formed by steep environmental gradients (Kark 2013). In terrestrial systems, species richness, abundances and productivity tend to peak in ecotones (Kark 2013, Smith et al 1997). However, the study of ecotones and its effects in the open ocean environment has not received the same attention as in terrestrial systems, probably because transition zones are not as easy to detect as in land. The best example of ecotones in the marine environment is oceanic fronts. Fronts are areas where distinct water masses meet creating enhanced horizontal gradients of physicochemical properties (e.g. temperature, salinity) (Belkin 2003). Fronts are widespread and linked to large effects on marine ecosystems (Le Fevre 1987, Longhurst 2006), including increased phytoplankton diversity (Barton et al 2010, Ribalet et al 2010) and productivity (Belkin et al 2009, Okkonen et al 2004, Springer et al 1996), yet their role in demarking bacterioplankton (i.e., defined as bacteria and archaea) communities shifts is unclear. This is critical since heterotrophic bacterioplankton make up the largest living biomass in the ocean, and drive oceanic biogeochemical cycles, regulating the composition of Earth’s atmosphere and influencing climate (Buchan et al 2014, Kirchman 2010). Prior work revealed oceanic fronts can act as ecotones, creating boundaries for bacterioplankton distribution in the ocean (Baltar et al 2016a). However, limited sampling and strong seasonal variability prevented testing of whether the front acted as a bacterioplankton diversity hotspot.

Here we performed the first seasonal high spatial resolution transect study of bacterioplankton diversity across an oceanic front (the Subtropical Frontal Zone off New Zealand). This study involved 6 sampling cruises from January 2014 to April 2015. We combined a high-resolution characterization of the front (via a continuous temperature and salinity recorder) and the analysis of bacterioplankton diversity (based on 16S rRNA gene Illumina sequencing (see Supplementary Information)). In all the cruises, bacterioplankton diversity and chlorophyll-a concentration was sampled at eight surface (2 m) water stations along the 48 km long transect, following the same approach as in previous studies (Baltar et al 2015, Baltar et al 2016a). The Subtropical Front is where the warm and salty subtropical waters (STW) and the cold, high-nutrient low-chlorophyll sub-Antarctic waters (SAW) meet. This front is constrained along the SE continental shelf of New Zealand to about 40-50 km offshore, compacting the frontal zone to a width of 2-10 km (Heath 1972).

Temperature and salinity changes along the transect retained the same structure throughout the study, with the presence of the 3 main water masses located in this area (Fig. 1A). Off the coast, the low salinity neritic waters (NW; characterized by riverine inputs) encounter and gradually mix with the warmer and saltier subtropical waters (STW). This is followed by a sharp (taking place in < 2 km between stations 3 and 5) drop in temperature and salinity, demarking the front and the convergence of the STW with the offshore SAW. As previously reported in other studies across this transect (Jillett, 1969; Heath, 1972; Shaw and Vennell, 2001; Hopkins et al., 2010), the location, width and strength of the front oscillates seasonally (Fig. 1B) (Heath 1972, Hopkins et al 2010, Jones et al 2013).

**Figure 1.**
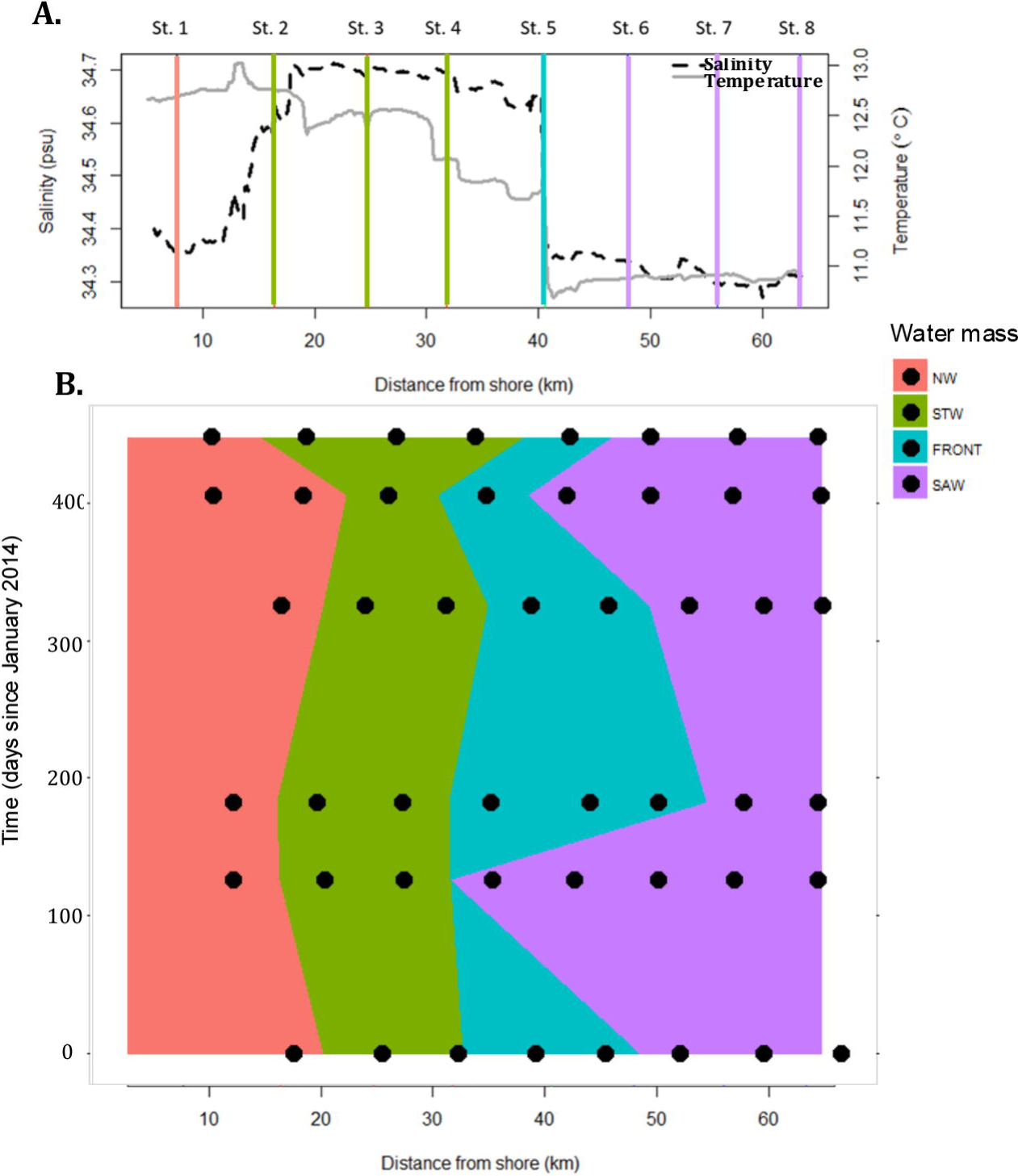
**(A)** Physicochemical changes in salinity (dashed) and temperature (solid) along the surface waters of the transect in April 2015, as a representation of the general structure of the water masses encountered in study region. The stations are shown as vertical lines, with their predicted water masses given by colour. Salmon pink = Neritic Water (NW), Green = Subtropical Water (STW), Blue = Frontal Water, Purple = Sub-Antarctic Water (SAW). This is an example of how the water masses were in April 2015, but their location shift throughout the study. **(B)** Summary of how the water masses shifted throughout the duration of the study, with sampling locations shown in black.

A clear water mass dependent seasonal change in phytoplankton biomass (chlorophyll-a) was observed along the transect (Fig. 2A). These blooms within the STW coincided with the transition from the Austral winter (June-July 2014) to the spring-summer seasons, as well as a smaller autumnal bloom. Observed chlorophyll-a concentrations reached >4 mg m^-3^ in December 2014 and >2 mg m^-3^ in April 2015 during the study, in agreement with previous studies where the threshold of >1 mg m^-3^ indicated bloom conditions (Hopkins et al 2010, Jones et al 2013). While strong seasonality in phytoplankton biomass was detected in the STW, changes in the NW and front were less pronounced, and absent within the SAW. This is consistent with the characteristic iron-limited primary production of the high-nutrient low-chlorophyll SAW (Baltar et al 2015, Jones et al 2013, Sander et al 2014). However, chlorophyll-a concentrations where not higher within the frontal waters at any time point, with values intermediate to those in STW and SAW, reflecting the mixing nature of fronts.

**Figure 2.**
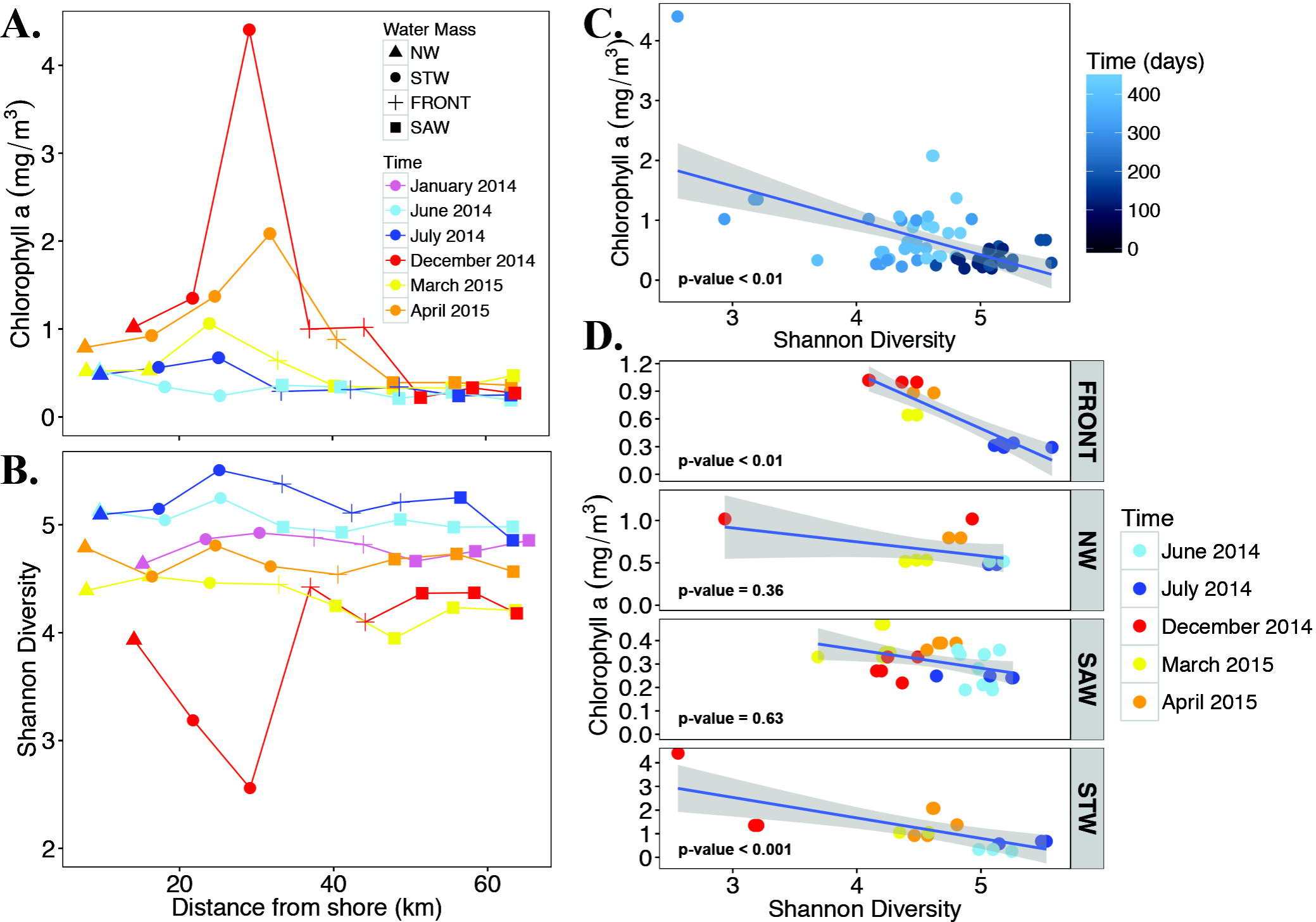
**(A)** Chlorophyll-a concentration, and **(B)** bacterioplankton diversity (Shannon index) in the surface waters along the transect in the six cruises performed from January 2014 to April 2015. Different sampling cruises are denoted in different colors, and the symbols denote different water masses. **(C)** Relation between chlorophyll-a concentration and bacterioplankton diversity (Shannon index) with all samples pooled together or **(D)** with samples separated by water mass. Neritic Water (NW), Sub-Antarctic Water (SAW), Subtropical Water (STW), frontal zone (FRONT).

Changes in chlorophyll-a concentration were negatively correlated (p < 0.01) to bacterioplankton diversity (based on the Shannon Index), with fluctuations across the transect modified by seasons (Fig2 B-D, and Tables S1-2). Changes in Shannon diversity were strongest across time (ANOVA F Ratio = 51.60, p value <.0001), with no significant effect across the water masses (p = 0.45) unless time was accounted for (water mass x time: F Ratio = 5.13, p value <.0001). While changes in bacterioplankton richness were observed and mirrored trends in diversity, it suggests that community structure (changes in evenness) where the strongest factor. Maximum diversity occurred during winter (June and July 2014) and the lowest in summer (December 2014). This inverse relationship between bacterioplankton diversity and phytoplankton biomass is consistent with the common negative diversity–productivity relationship found in aquatic ecosystems (Baltar et al 2016b, Smith 2007). This is specifically pronounced in areas with larger blooms (STW and the frontal waters) (Fig. 2D), which are the water masses with the strongest seasonal variations in chlorophyll-a and bacterioplankton diversity. Consistent with observations for phytoplankton biomass, bacterioplankton diversity did not peak in the front, with diversity values intermediate to those for SAW and STW.

Our results demonstrate that while fronts serve as ecotones in the sense of delimiting the distribution of bacterioplankton, their strength is not constant, but seasonally driven. This seasonality is linked to differences in phytoplankton biomass across the different water masses (e.g. December 2014). While oceanic fronts are clear boundaries, we demonstrate that they are not bacterioplankton diversity hotspots in contrast to observations for terrestrial plants. This might be due to the more dynamic nature of oceanic fronts, that experience continuous mixing, enlarging and contracting events resulting in a less stable environment than terrestrial ecotones.

## Acknowledgements

We would like to thank the skippers and crew of the RV Polaris II for their help during the sampling events. FB was supported by a University of Otago Research Grant (UORG).

### Conflict of interest

The authors declare no conflict of interest.

